# Genetic analysis of DinG-family helicase YoaA and its interaction with replication clamp-loader protein HolC in *E. coli*

**DOI:** 10.1101/2021.01.04.425237

**Authors:** Vincent A. Sutera, Thalia H. Sass, Scott E. Leonard, Lingling Wu, David J. Glass, Gabriela G. Giordano, Yonatan Zur, Susan T. Lovett

## Abstract

The XP-D/DinG family of DNA helicases contribute to genomic stability in all three domains of life. We investigate here the role of one of these proteins,YoaA, of *Escherichia coli*. In *E. coli*,YoaA aids tolerance to the nucleoside azidothymidine (AZT), a DNA replication inhibitor and physically interacts with a subunit of the DNA polymerase III holoenzyme, HolC. We map the residues of YoaA required for HolC interaction to its C-terminus by yeast two-hybrid analysis. We propose that this interaction competes with HolC’s interaction with HolD and the rest of the replisome;YoaA indeed inhibits growth when overexpressed, dependent on this interaction region. By gene fusions we show YoaA is repressed by LexA and induced in response to DNA damage as part of the SOS response. Induction of YoaA by AZT is biphasic with an immediate response after treatment and a slower response that peaks in the late log phase of growth. This growth-phase dependent induction by AZT is not blocked by *lexA3* (Ind^-^), which normally negates its self-cleavage, implying another means to induce the DNA damage response that responds to the nutritional state of the cell. We propose that YoaA helicase activity increases access to the 3’ nascent strand during replication; consistent with this,YoaA appears to aid removal of potential A-to-T transversion mutations in *ndk* mutants, which are prone to nucleotide misincorporation. YoaA and its paralog DinG also may initiate template-switching that leads to deletions between tandem repeats in DNA.

**IMPORTANCE:** Maintaining genomic stability is crucial for all living organisms. Replication of DNA frequently encounters barriers that must be removed to complete genome duplication. Balancing DNA synthesis with its repair is critical and not entirely understood at a mechanistic level.The YoaA protein, studied here, is required for certain types of DNA repair and interacts in an alternative manner with proteins that catalyze DNA replication. YoaA is part of the well-studied LexA-regulated response to DNA damage, the SOS response. We describe an unusual feature of its regulation that promotes induction after DNA damage as the culture begins to experience starvation. Replication fork repair integrates both DNA damage and nutritional signals. We also show that YoaA affects genomic stability.

## INTRODUCTION

The YoaA protein of *Escherichia coli* is a member of the XP-D/DinG family of DNA helicases, with members found in all three domains of life. These are superfamily 2 helicases with shared property of 5’ to 3’ translocation on ssDNA and an intrinsic Fe-S cluster. In humans, these proteins play various roles in DNA repair and the maintenance of genomic stability, the loss of which results in a variety of genetic diseases (1-4).

The bacterium *Escherichia coli* encodes two paralog proteins of this family, DinG and YoaA, both of which appear to be induced by DNA damage as part of the SOS response,(5-7). DinG encodes a structure-specific DNA helicase with the ability to unwind D-loops, R-loops and G-quadruplex sequences (8-10). Despite its induction by UV irradiation, *dinG* mutants show only a slight sensitivity to UV (8). Along with two other SF1 helicase proteins, UvrD and Rep, DinG appears to enhance survival of head-on replication/transcriptional collisions in vivo (11), when highly transcribed regions of the chromosome are inverted.

YoaA was identified in a genetic screen for factors that promote tolerance to the chain-terminating nucleoside azidothymidine (AZT) in *Escherichia coli* (12). AZT is incorporated during DNA replication and, since it blocks DNA chain elongation, produces single-strand DNA (ssDNA) gaps at the replication fork; cells can tolerate certain levels of AZT through its removal from DNA by exonuclease III (13) or DnaQ proofreading (12). Mutants in *yoaA* are viable but are strongly sensitive to AZT (12) as well as to MMS (14). Mutants in *dinG* are only very slightly AZT sensitive but do further enhance the sensitivity of *yoaA* mutants when combined.

YoaA is of particular interest because it physically interacts with the replisome protein, HolC (χ) of DNA polymerase III (12, 15, 16). Increased expression of HolC, like YoaA, promotes tolerance to AZT in vivo (12). HolC is purified as an intrinsic component of the DNA polymerase III, where it serves as an accessory protein to the clamp loader complex (17, 18). It is the one component of the replisome that interacts with single-strand DNA binding protein, SSB (19). In addition to its interaction with SSB, HolC forms an heterodimeric complex with HolD (ψ); it is HolD that links this accessory dimer to the clamp loader and to the rest of the replisome (20, 21).

The HolC/YoaA complex and the HolC/HolD complex appear to be mutually exclusive structures (22). The same residues buried at the HolC/HolD interface, F64 and W57 (23), are also required to form a complex with YoaA and essential for AZT tolerance in vivo (22). Expression of YoaA/HolC/HolD yields two complexes, HolC/HolD and HolC/YoaA, with no evidence of a ternary complex (22). This finding led to the hypothesis that HolC forms two complexes, HolC/HolD dedicated to replication and HolC/YoaA to repair, both recruited to ssDNA through HolC’s SSB interaction.

In this study we investigate further the genetic role of YoaA. Using yeast two-hybrid analysis, we map residues required for HolC interaction to the C-terminal 18 amino acids of YoaA and show that they are required for YoaA function in vivo. When strongly overexpressed and its C-terminus is intact,YoaA inhibits growth. By gene fusions, we confirm LexA repression of YoaA expression and its induction by AZT; mutation of the putative LexA box at 24 nucleotides upstream of the open reading frame yields constitutively high expression even in the absence of damage. A non-inducible allele of LexA, *lexA3*, blocks the bulk of AZT-induction of *yoaA* expression, although there remains some residual induction of P*yoaA* by AZT, especially during the transition of the culture from exponential growth to stationary phase, suggesting an alternative mechanism for overcoming LexA repression at the locus, induced by starvation. Consistent with our hypothesis that YoaA provides access to the 3’ nascent strand during replication, elevated YoaA expression promotes template-switching that produces deletions between DNA tandem repeats and reduces T- to A transversion mutations in a *ndk* mutant prone to nucleotide misincorporation.

## Results

### Determination of HolC binding residues within YoaA

The alignment of *Escherichia coli* DinG and YoaA proteins (Figure 1) shows 29% identity over the length of the two proteins, including all helicase motifs (Q motif, I, Motifs Ia, II, III, IV,V,VI, and P motif), the two helicase HD1 and HD2 domains, and the 4 cysteine residues that coordinate Fe-S binding (24). In our prior study, we showed that K51(Walker A, motif I), C168 (Fe-S cluster) and D225 (Walker B, motif II) (see Figure 1) were required for *yoaA* to promote AZT tolerance in vivo (12). The most diverged regions of the YoaA and DinG proteins are the Arch domain (between Motifs II and III) and the C-terminus. A clue to the YoaA region that binds HolC came from prior pulldown experiments, where we noted that a C-terminally truncated proteolytic fragment of YoaA present in the extracts failed to pulldown with HolC as did full-length YoaA (12).

**Figure 1.**
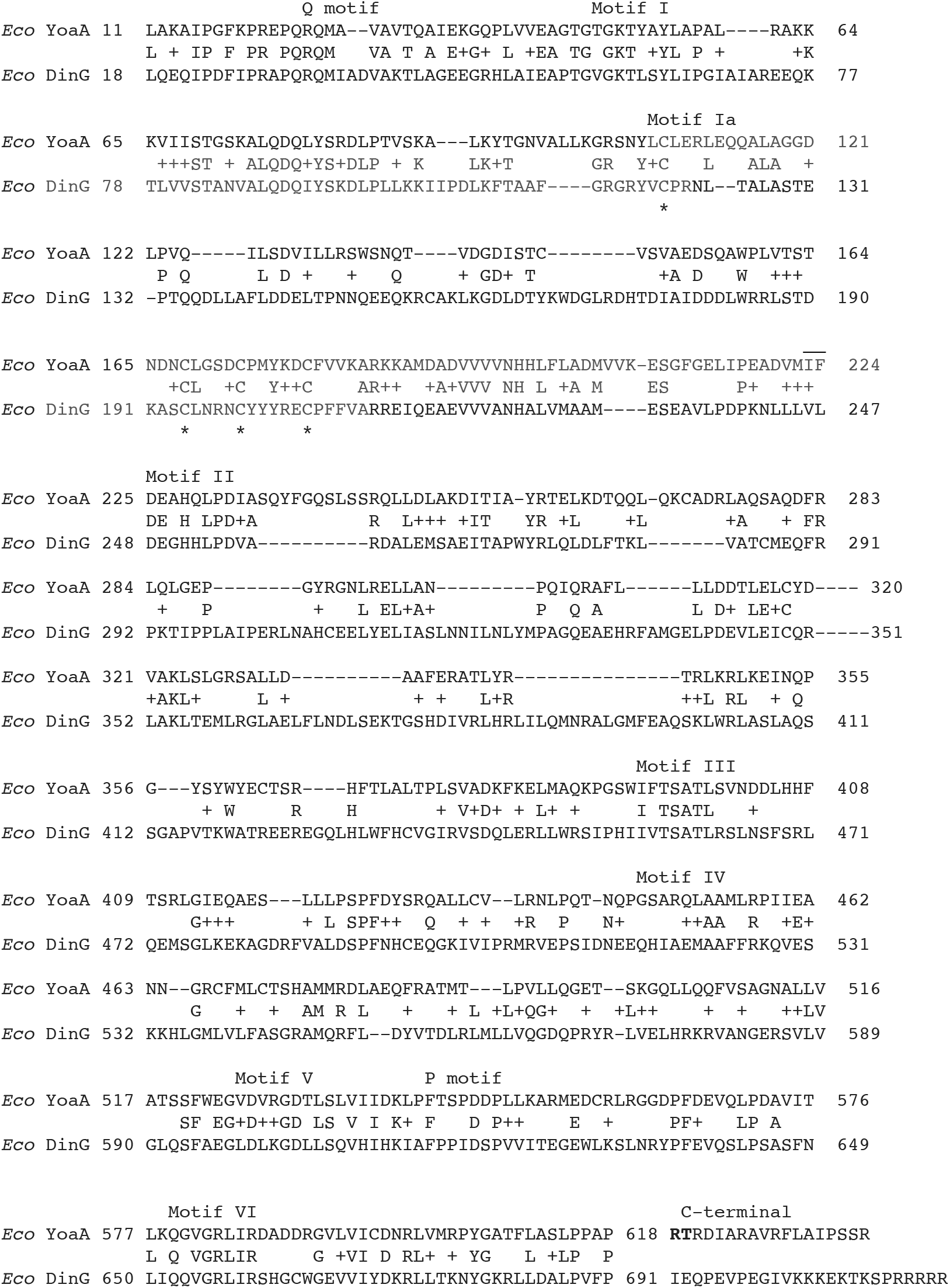
BLAST Alignment of YoaA and DinG proteins. Conserved helicase motifs are indicated above and cysteine residues of the FeS cluster are marked below with an asterisk. YoaA R619 and T620, implicated in HolC binding, are shown in bold at the C-terminus.

We deleted the C-terminal amino acids of YoaA on a plasmid-expressed His_6_-tagged allele and assayed the ability of it to complement the AZT-sensitivity phenotype of *yoaA* mutants (Figure 2).

**Figure 2.**
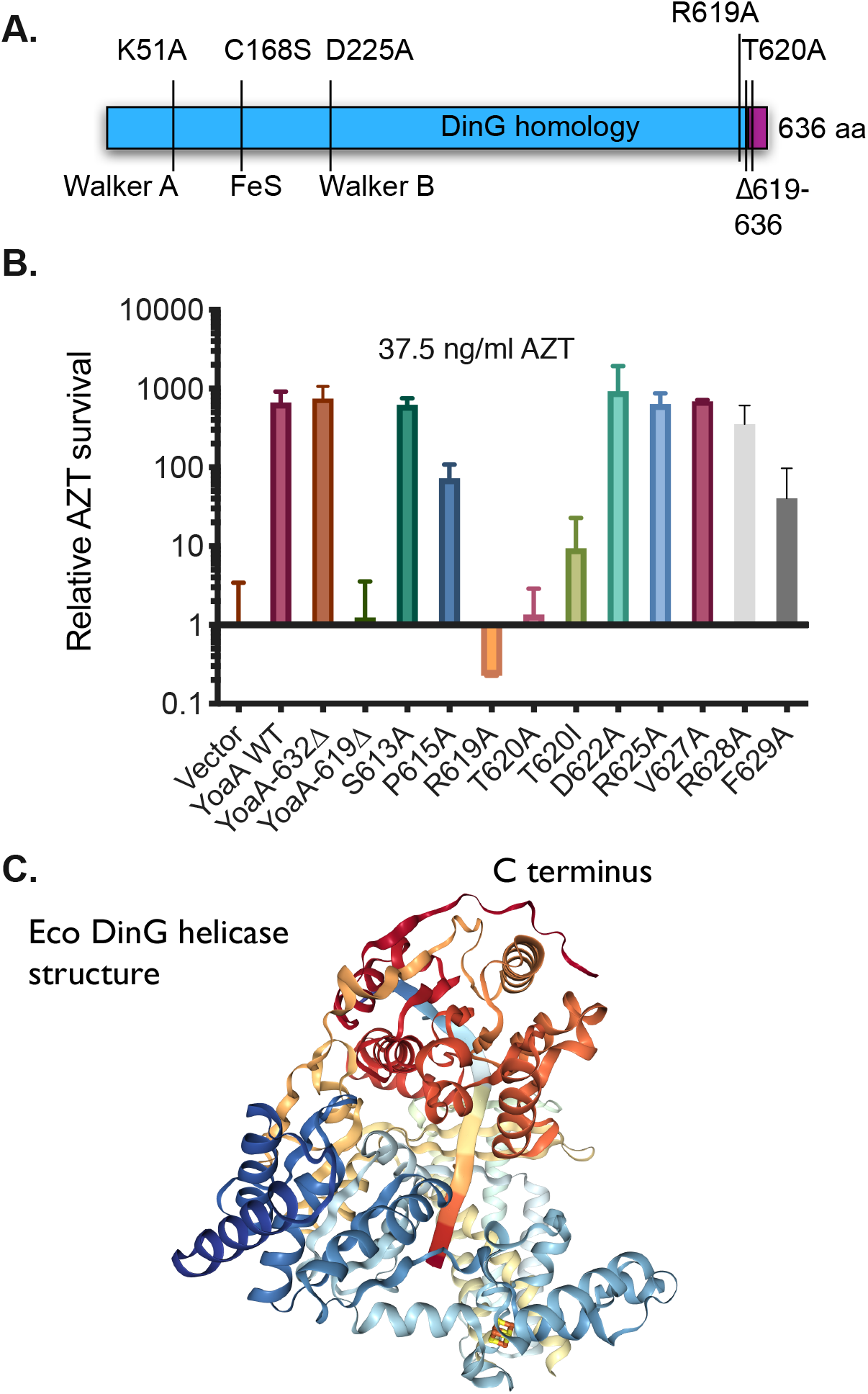
YoaA complementation assays. A. Schematic of the *yoaA* gene showing previously identified non-complementing mutations (Brown et al. 2015) and those identified in this study. The C-terminal region dissimilar to Eco DinG is denoted in purple. B. Complementation assay. Plating efficiency at 37.5 ng/ml AZT of strains carrying the designated *yoaA* plasmid alleles in a *yoaA*Δ strain relative to the vector control. Fractional survival at this dose was 0.66 for *yoaA*^+^ and 0.0009 for the vector control. Error bars represent standard deviations. C. Eco DinG structure. Image from the RCSB PDB (rcsb.org) of PDB ID 6FWR (Cheng and Wigley 2018), with the C-terminus indicated. DNA within the structure is shown as a flat ribbon.

(Although the *yoaA* gene is transcribed from the *tac* promoter on these plasmids, we were able to detect complementation without the addition of inducer IPTG.) Whereas the wild-type allele and *yoaA*Δ632-636 enhanced survival at 37.5 ng/ ml dose of AZT, almost 1000-fold relative to the plasmid vector control, the *yoaA*Δ619-636 allele failed to complement, with a plating efficiency on the AZT medium similar to the vector control. We mutated a number of individual residues within the region between residues 619 and 631 and found that both R619A and T620A destroyed YoaA function as measured by AZT tolerance, whereas D622A, R625A,V627A, and R628A had no effect; F629A and T620I showed a partial loss of complementation.

After inducing plasmid expression with IPTG,Western blotting of biotin-binding domain tagged YoaA,YoaAΔ619-636,YoaA R619A and YoaA T620A showed levels of soluble YoaA protein comparable to wild-type, indicating that failure to complement was not a result of protein degradation (Figure 3). We confirmed that YoaAΔ619-636,YoaA R619A and YoaA T620A conferred AZT-sensitivity when transferred to the native, naturally expressed *yoaA* locus on the *E. coli* chromosome, showing that the dysfunction of these alleles is not plasmid-dependent. (Supplemental Figure 1).

**Figure 3.**
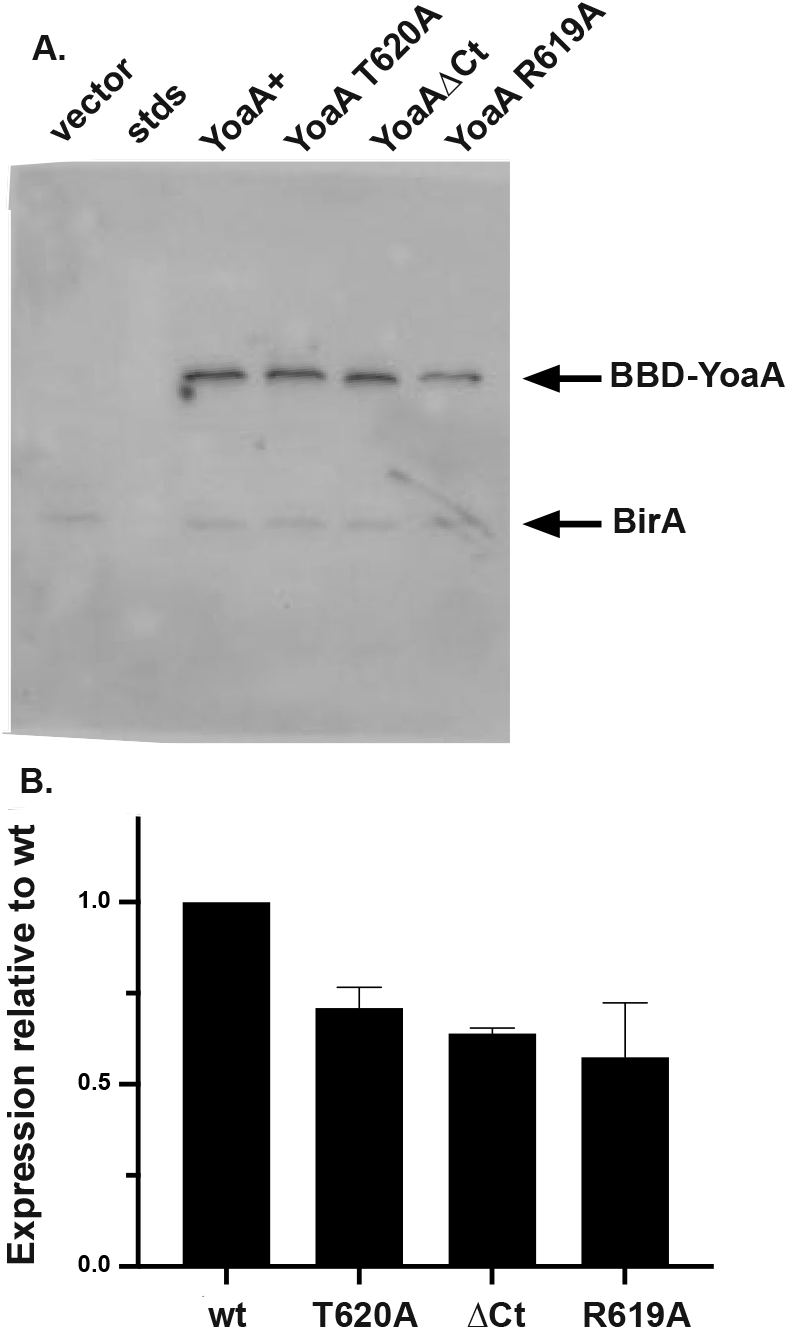
Western blot of BBD-fused YoaA carrying the indicated alleles (“ΔCt”= Δ619-636 of YoaA) using Neutravidin detection, compared to cells expressing vector. BirA is biotin binding protein of E. coli. A. Representative gel. B. Quantitation of Western blots, showing average and range of 2 independent experiments.

Based on its similarity with DinG for which there is structural information, the location of the C terminus of the YoaA is likely to be on the outside of the protein, at a site distinct from those involved in ATP and DNA binding. The corresponding C-terminus of DinG is partially visible in crystal structures (PDB ID 6FWS, 6FWR) on the exterior surface of the HD2 domain where it overlies helix 18, containing helicase motif IV. (Figure 2C. Image from the RCSB PDB (rcsb.org) of PDB ID 6FWR (24).)

We transferred the most defective *yoaA* alleles to yeast two-hybrid fusions to ascertain whether these YoaA alleles retain the ability to interact with HolC, as we have demonstrated previously (12, 16). Whereas wt YoaA showed an interaction with HolC, indicated by growth on the -His plates, YoaA R619A,T620A and the YoaAΔ619-636 C terminal truncation did not. All strains grew equally well on -Trp Leu plates, which select for the presence of the two plasmids. Control plating of individual plasmids combined with a vector control partner were performed in parallel and yielded negative results (data not shown) so the interaction requires both HolC and YoaA fusion partners. We obtained similar results whether the activation domain was fused to either HolC or YoaA, with the corresponding partner fused to the DNA binding domain, in three independent plating experiments (Figure 4).

**Figure 4.**
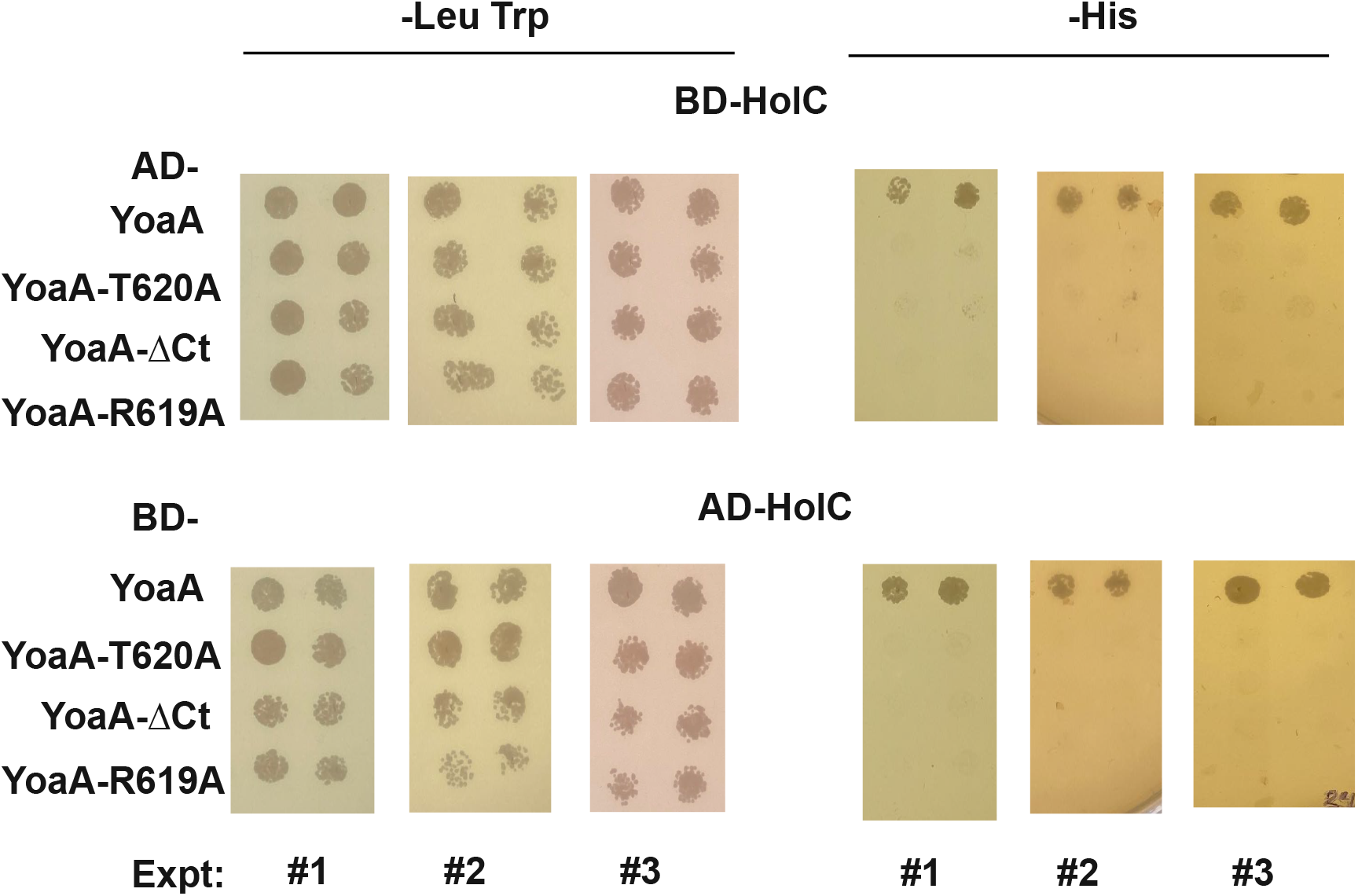
Yeast two-hybrid analysis of YoaA and HolC interaction. Shown at the left are segments of -Leu Trp plates that select for the two hybrid plasmids; at right are those from - His plates that selection for a functional interaction. The top row shows results from hybrids in which the HolC is fused to the Gal4 DNA binding domain (BD) and YoaA (wt and three mutants alleles) is fused to the Gal4 activation domain (AD). YoaAΔCt is the YoaA truncation at amino acid 618, YoaAΔ619-636. In the bottom series HolC is fused to the activation domain and YoaA and its alleles are fused to the DNA binding domain. Three independent experiments are shown.

### Excess YoaA is toxic, dependent on an intact C-terminus

HolC binds to HolD, a second protein in the clamp-loader complex of DNA pol III, and both proteins are required to sustain full viability and fast growth (25). Because YoaA also forms a complex with HolC, as an alternative to that of HolC/HolD (22), excess YoaA is expected to interfere with growth, by competing with HolD for HolC. Induced expression of YoaA from a high copy number plasmid driven by the *tac* promoter was indeed found to inhibit the growth of otherwise wild-type strains (Figure 5), whereas uninduced cultures grew unperturbed. A YoaA derivative lacking the C-terminal 18 amino acids required for interaction with HolC was not toxic, nor was the K51R mutation of the Walker A box.

**Figure 5.**
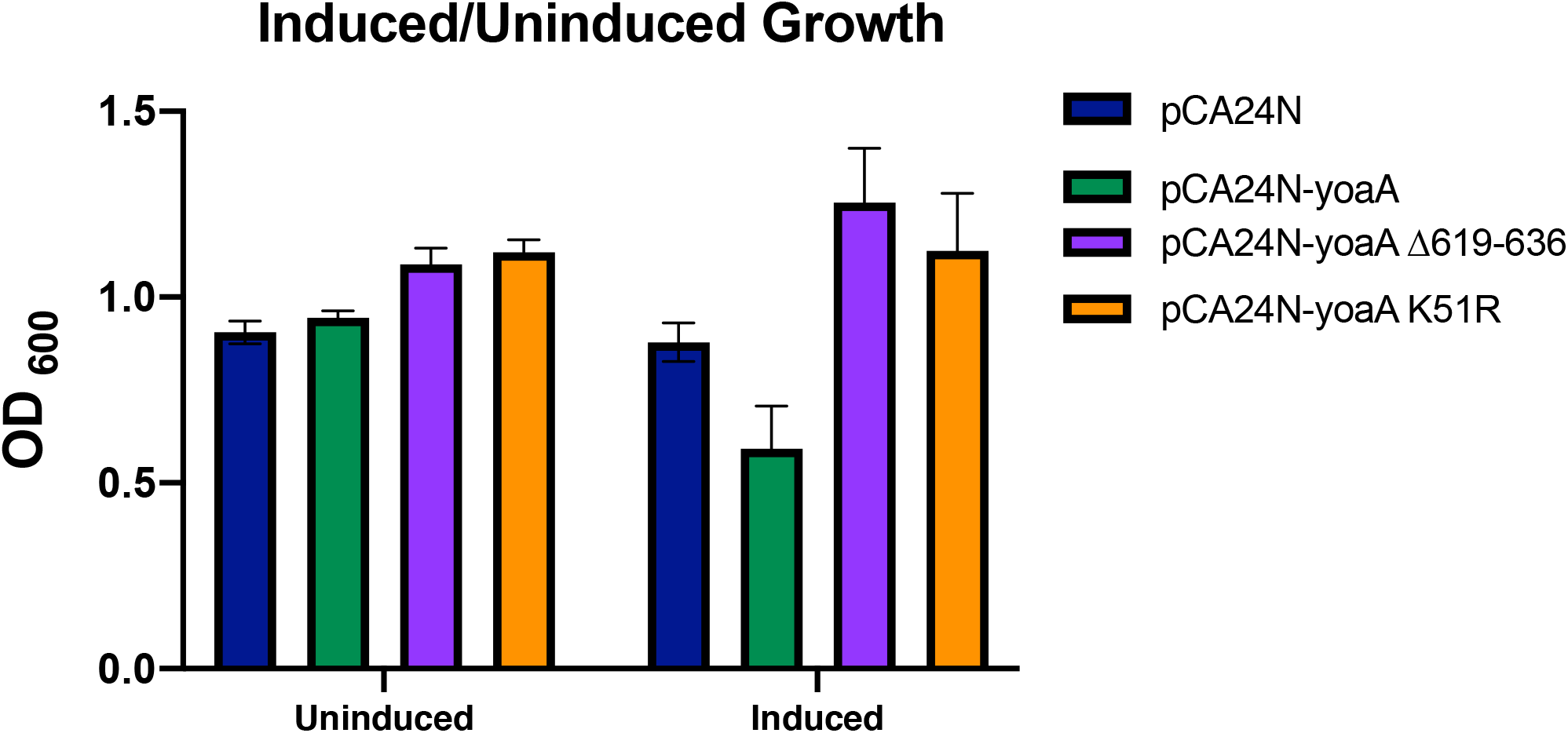
Growth inhibition by expression from pCA24N-YoaA plasmids, with cultures split and then grown in LB (left) or LB + IPTG (right). Plotted are the final mean ODs of 6 independent cultures 3 h after the addition of IPTG, with error bars indicating standard deviations.

### Expression of YoaA

To study the regulation of the *yoaA* gene, we fused its upstream region (intergenic with divergently-transcribed *yoaB*, with a length of 132 bp) to the *Photorhabdus luminescens* luciferase *luxCDABE* operon on a plasmid and measured luminescence and OD through the growth of the culture, with and without AZT addition (to 1.25 ng/mL) at time zero. This is a sublethal concentration of AZT for wt strains, where cells continue to proliferate. Expression was induced rapidly after addition of AZT to the culture, with a peak approximately 60 minutes after treatment, as we have observed for other similar fusions to SOS-regulated promoter regions of *recA* and *dinB* (13). In contrast to those fusions, for *yoaA::lux* we consistently saw a biphasic induction curve, with a slower second induction in late phases of growth, reaching a maximum approximately 150-180 minutes after treatment. A LexA mutation, *lexA3*, affecting the proteolytic cleavage site that renders the SOS response non-inducible, negates most induction of expression by AZT, especially that soon after treatment, although the second slow phase of induction remained partially intact (Figure 5A). At 180 min after AZT addition, expression in the *lexA3* genetic background is approximately 30% that of wt and 6-fold higher in treated vs. untreated cells. Note that the data are normalized to culture OD, so the slow increase in expression is not due to expansion of the culture. In a second experiment we compared the induction of *yoaA* expression by AZT from an intact upstream region with one in which we mutated the putative LexA box (replacing the two triplets of invariant LexA box sequence CTG(N)_10_CAG with CCC(N)_10_GGG; (N)_10_ = ttcaaatcaa for *yoaA*). As expected for a LexA-repressed gene, we saw high constitutive expression in the absence of any damage, with no further increase by the addition of AZT (Figure 5B).

### YoaA does not affect recombination

Because a number of eukaryotic members of the XPD/DinG helicase family affect homologous recombination or mutation rates, we examined whether loss of *yoaA* had effects on several assays for recombination or genetic instability (Figure 6). We measured crossover recombination using an assay previously developed that varies the amount of homology between the recombining loci (26). Recombination at limiting homology often reveals genetic effects not apparent with larger homologies (26, 27). This assay primarily measures the RecFOR pathway of homologous recombination, which is believed to be the primary pathway in *E. coli* for recombination at single-strand gaps in DNA (reviewed in 28). We saw no influence of *yoaA* at any homology length using this crossover assay (Figure 7A).

**Figure 6.**
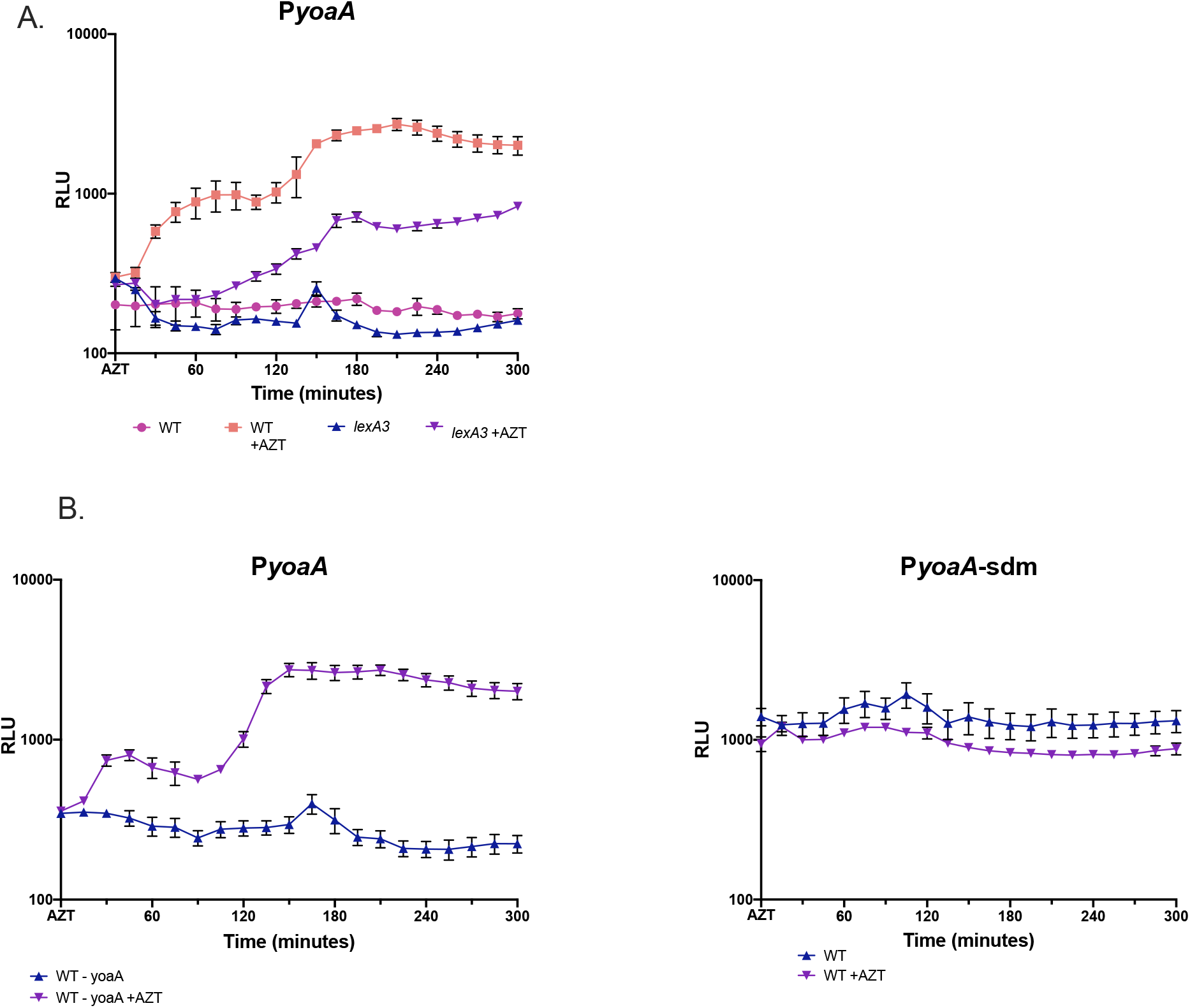
YoaA promoter (P*yoaA*) expression as measured by *lux* operon fusions during growth of the culture in LB, with and without addition of AZT at time 0. Values are expressed as relative luminescence units, with luminescence cpm divided by OD_600_ of the culture at that time. The average of 4 replicates is plotted, with errors indicating standard error of the mean. A. Expression from the *yoaA* 132 bp upstream intergenic region(in wt strains, with and without AZT compared to *lexA3* (noninducible), with and without AZT. B. Left panel: expression of the *yoaA* upstream intergenic region from wt *yoaA* in wt strains with and without AZT; right panel and performed in parallel with the left: expression of the promoter region with a site-directed mutation, P*yoaA*-sdm to remove the putative LexA box at -24, with and without AZT.

**Figure 7.**
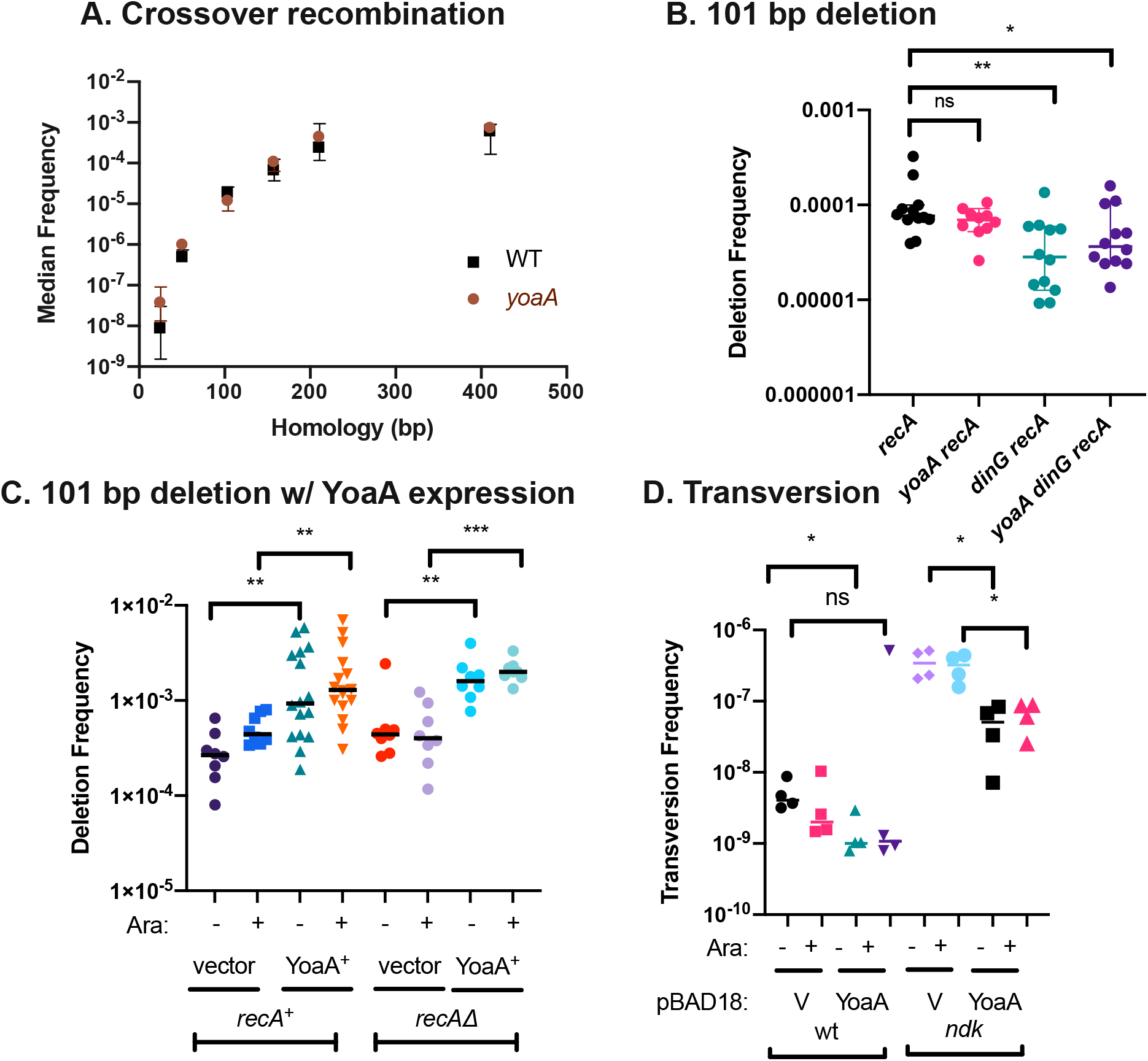
A. Recombination frequencies in wt and *yoaA* strains at differing amounts of homology. B. RecA-independent deletion frequencies between 101 bp tandem repeats in mutants of *dinG* and/or *yoaA*. Bars indicate median values and the Mann-Whitney significance levels indicated with astericks. The deletion assay plasmid is pSTL57. C. Deletion frequencies of wt or *recA*Δ carrying either the pBAD18 vector or a BAD18 plasmid carrying *yoaA*^+^. Cultures were split and treated for 2 hours with 0.2% arabinose (+) or not (-). The deletion assay plasmid is pSTL141. Bars indicate median values and the Mann-Whitney significance levels indicated with astericks. D. A to T transversion frequencies in wt or *ndk* mutant strains, carrying either the pBAD18 vector (V) or *yoaA*, with and without arabinose induction.

### YoaA and its paralog DinG promote template-switching and genomic rearrangements

Tandem direct repeats in DNA are unstable and prone to deletion. In *E. coli* deletion between 101 bp tandem repeats occurs at high frequency in the population during DNA replication, independent of the homologous recombination factors, including RecA (reviewed in 29). Previous work supports a template-switching model for rearrangements between tandem repeats involving misalignment of the nascent strand.

Our model of how YoaA may facilitate replication fork repair (12) is that its 5’ to 3’ helicase activity may unwind the the nascent 3’ terminus, allowing repair factors increased access to it (Figure 8). In the template-switching model for deletion formation (reviewed in 29), a nascent 3’ terminus is unwound and mispairs with a second copy of the repeat, either on its downstream template or across the fork to the sister nascent strand (Figure 8). It was therefore of interest to determine if YoaA or its paralog DinG had any effects on deletion frequencies between tandem repeats since they are candidates for the functions that could initiate a template-switch. A knockout of *yoaA* did not affect RecA-independent deletion frequencies measured between 101 bp repeats in the *tetA* gene of plasmid pBR322 (Figure 7B). However, we saw a modest but significant reduction of deletion frequencies 2-3 fold in both *dinG* single and *dinG yoaA* double mutants, suggesting that DinG promotes some spontaneous deletion events measured in this assay (Figure 7B). The lack of effect by *yoaA* does not necessarily mean it does not stimulate deletion formation—its effects may be hidden by a larger contribution by DinG. To test whether YoaA could promote deletion when its expression was elevated, we cloned the gene under the control of a pBAD arabinose promoter. Arabinose-induced expression of YoaA from this pBAD18 plasmid stimulated deletion of 101 bp *tetA* repeats about 5-fold, compared to control strains carrying the pBAD18 vector (Figure 7C). We saw a substantial but smaller stimulation of deletion by *yoaA* even without induction of the *ara* promoter. We were unable to test effects of DinG expression similarly due to its toxicity.

**Figure 8.**
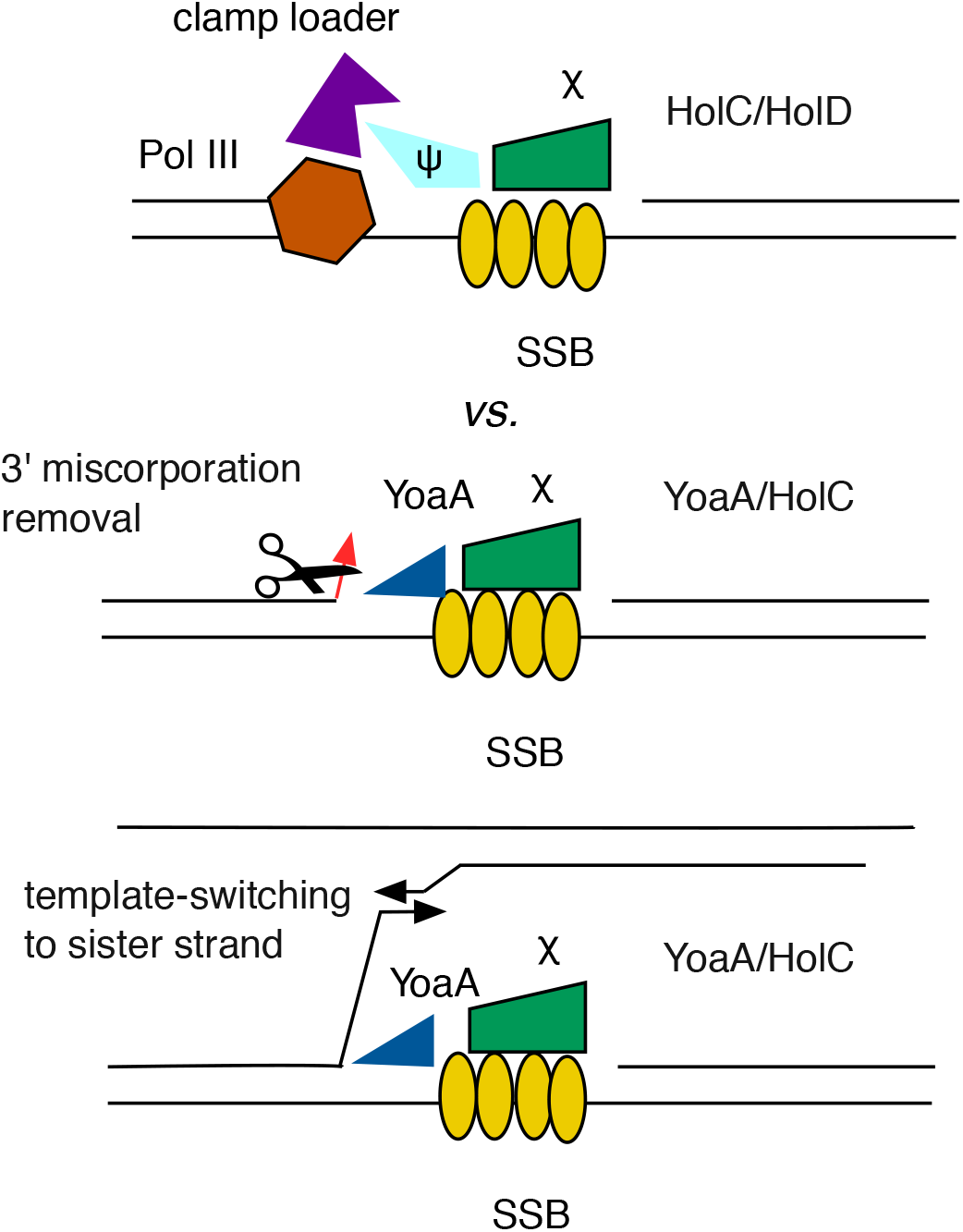
Diagram of HolC complexes and reactions they provoke. Top: Through its interaction with SSB, HolC recruits HolD and the clamp loader and DNA pol III replisome. Middle: Alternatively, through its interaction with SSB HolC recruits YoaA helicase, which unwinds the 3’ nascent strand, allowing exonucleases access for removal of terminal misincorporation or AZT. Bottom, recruitment and unwinding by YoaA through HolC may also initiate a template-switch to the sister strand, producing repeat rearrangements and sister chromosome exchange.

### YoaA may enhance proofreading

If YoaA enhances access to the 3’ nascent strain, it may assist in the removal of incorrect nucleotides incorporated during replication, either by intrinsic (DnaQ-dependent) or extrinsic (DnaQ-independent) proofreading. Mutants in nucleotide diphosphate kinase, *ndk*, exhibit a mutator phenotype due to misincorporation caused by perturbations of dNTP pools (30-32). A to T transversions, in particular, are elevated in *ndk* mutants; these are not affected by mismatch repair proficiency but are strongly influenced by proofreading by the DnaQ (ε) subunit of Pol III (31-33). Using specific *lacZ* revision assays, we examined effects of YoaA expression on A to T transversions in wt and *ndk* mutants. We observed a 120-fold stimulation of A to T transversion frequencies by loss of *ndk* (Figure 7D). Plasmid-expressed YoaA^+^, decreased transversion frequencies about 5-7 fold in *ndk* mutants. We saw a smaller reduction of transversion frequencies, 2-4 fold, by YoaA expression in wt strains. As we had seen previously for *yoaA* expression effects on deletion frequencies, induction of the pBAD promoter with arabinose had little effect.

## DISCUSSION

In this study we examine further the genetic effects of the *yoaA* gene of *E. coli*. We mapped the residues of YoaA required for HolC binding to the C-terminal 18 amino acids of the protein, a region in which it is distinct from its paralog protein, DinG. Mutation of two specific residues R619 and T620 to alanine also abolished interaction with HolC, as determined with yeast two-hybrid analysis. Mutants in *yoaA* are sensitive to the replication inhibitor AZT; the ability of YoaA to interact with replisome protein HolC is likely required for AZT tolerance, since YoaA R619A and T620A and C terminal deletions fail to complement this defect and cause AZT sensitivity when these alleles are introduced into the natural *yoaA* gene on the chromosome. These mutations had only minor effects on YoaA protein accumulation in vivo and, based on similarity of the protein to the DinG helicase for which there is structural information (24), likely affect a peripheral region of the helicase. Whether this interaction is merely a means for recruitment of the helicase to persistent ssDNA gaps, through a HolC/SSB interaction, or in someway alters the properties of the enzyme remain to be determined.

A knock-out mutation of *yoaA* does not affect normal growth whereas a knock-out mutation of *holC* or *holD* produces slow growth and inviability, especially on rich media (25, 34). If YoaA/ HolC forms an alternative complex to HolC/HolD as we have proposed (22), increased expression of YoaA may lead to competition and a slow growth phenotype by interfering with the formation of the growth-promoting HolC/HolD complex. Expression of YoaA from a strong promoter on a high-copy number plasmid does indeed inhibit growth whereas expression of C-terminally truncated YoaA does not. Therefore, the control of YoaA expression is likely critical to balancing repair vs. replication functions.

We confirm that *yoaA* is a DNA-damage inducible gene by promoter fusions to a luciferase (*luxCDABE*) operon, as suggested by previous microarray experiments, with and without UV exposure (7). We also demonstrate LexA-regulation of the gene, with a non-inducible *lexA3* allele reducing AZT induction and mutation of the predicted LexA box causing constitutively high expression. One advantage of *luxCDABE* luciferase fusions is that the expression signal can be detected in live cells (because the substrate is an endogenous metabolite), allowing us to follow expression during growth of the culture in the presence of a sublethal concentration of AZT. Following AZT treatment of an early log phase culture, *yoaA* shows an unusual bimodal induction of expression, with the first increase within 60 minutes, as is typical of SOS genes (see 13)). AZT induces replication gaps and is a strong inducer of the SOS response via the RecAFOR pathway (13) that promotes the cleavage of the LexA repressor. In addition to the immediate response, we observed a slower secondary increase as the culture begins to approach stationary phase. Note that because AZT is incorporated during replication, it would not be expected to affect stationary phase, non-replicating cells. The *lexA3* mutation, which prevents its cleavage (35), abolishes the immediate mode of induction, but does not negate the slow and more gradual induction at late phases of growth. We hypothesize that there is nutritional modulation of LexA repression of the *yoaA* gene, such that induction of the gene is triggered more easily as the culture ages. We suspect that this is advantageous because the last rounds of replication may become more difficult to complete as cells begin to starve. Because this induction persists in the non-cleavable *lexA3* strain, there may be an alternative mechanism to relieve LexA gene repression in addition to RecA-stimulated self-cleavage. To our knowledge, this phenomenon has not been reported previously, although most studies have not been conducted with late log phase cultures. The generality and the mechanism of this second phase of induction therefore remains be be determined. The regulation of YoaA is likely important because of its competition with HolD and the replisome for interaction with HolC/SSB and YoaA expression may therefore interfere with replication.

Although YoaA promotes tolerance of ssDNA gaps caused by AZT incorporation in DNA, we do not find that it alters the efficiency of the RecAFOR pathway of homologous recombination during normal growth. YoaA had both positive and negative effects on genetic stability. Elevated expression of YoaA reduced A to T transversion mutations in a nucleotide diphosphate kinase *ndk* mutant that is prone to misincorporation during replication due to perturbed dNTP pools.

This observation is consistent with YoaA promoting proofreading, either intrinsic or extrinsic to Pol III, following nucleotide misincorporation. Conversely, elevated expression of YoaA promoted template-switching that leads to RecA-independent deletions formed between 101 bp tandem repeats. When we discovered that RecA-independent rearrangements were accompanied by crossing-over between sister chromosomes, we proposed that this pathway could function during replication gap repair and may be initiated by a DNA helicase (36). The data presented here suggests that both YoaA and DinG helicases are candidates for this role and that both can provoke template-switching during replication. DinG appears to play a larger role during normal growth, since its loss reduces constitutive levels of rearrangements, whereas YoaA loss does not but affects deletion rates when its expression is elevated. Both YoaA and DinG promote tolerance to AZT, but YoaA plays the larger role of the two. How these paralog proteins are specialized and their cellular roles remains to be explored more fully by further genetic and biochemical characterization.

## MATERIALS AND METHODS

### Strains, plasmids, growth conditions

For this study we used *Escherichia coli* K-12 strains MG1655 as wild-type (wt, *rph*-1) and isogenic strains STL3817 (*recA*::*cat*), STL9813 (*yoaA*Δ::FRT), STL9820 (*yoaA*Δ::FRT *recA*::*cat*), STL9824 (*dinG*::*kan recA::cat)*, STL9822 (*yoaA*Δ::FRT *dinG*::*kan recA*::*cat*), STL22918 (*lexA3 malF3089*::Tn10), STL13722 (*lacZ*-T1385A *mphC281*::Tn10), STL15952 (*lacZ*-T1385A *mphC281*::Tn10 *ndk*Δ::FRT *kan*. LB (37), Lennox formulation, was used for standard growth media. Plate media included the addition of Bacto-agar at 2%. For plasmid selection, the following antibiotics were employed at the given concentrations: ampicillin (Ap) at 100 µg/ml, kanamycin (Km) at 60 µg/ml, tetracycline (Tc) at 15 µg/ml, phleomycin (Phleo) at 5 µg/ml, and chloramphenicol (Cm) at 15 µg/ml. Strains were grown at 37°C. Minimal 0.2% lactose medium (38) was used to select Lac+ revertants in mutation assays.

For budding yeast, YEPD media (complete) and Drop Out Media (synthetically deficient) followed published recipes (39). Strains were incubated at 30°.

Site-directed mutagenesis of plasmid pCA24N-YoaA+ was used to create YoaA mutants at specific residues. Plasmids and primers are listed in Table 1. The forward primer was phosphorylated with T4 polynucleotide kinase (New England Biolabs) and used in a high fidelity PCR reaction (see below) with its complement primer. Following DpnI digestion (New England Biolabs, the PCR product was purified (BioBasic Inc.), ligated with T4 DNA ligase (New England Biolabs) and and transformed into the host strain XL1-Blue by electroporation (40). PCR reactions employed Pfu DNA polymerase from Agilent Technologies®, using the guidelines provided from the manufacturer. To create each *yoaA* site-directed mutant, primers that were used are listed in Table 1. Plasmids (listed in Table 1) from bacterial transformants were isolated using BioBasic® Inc. plasmid purification kits and the procedure of the manufacturer. DNA sequence analysis (GeneWiz) confirmed the presence of each particular *yoaA* site-directed mutant and no other change to the sequence.

**Table 1.**
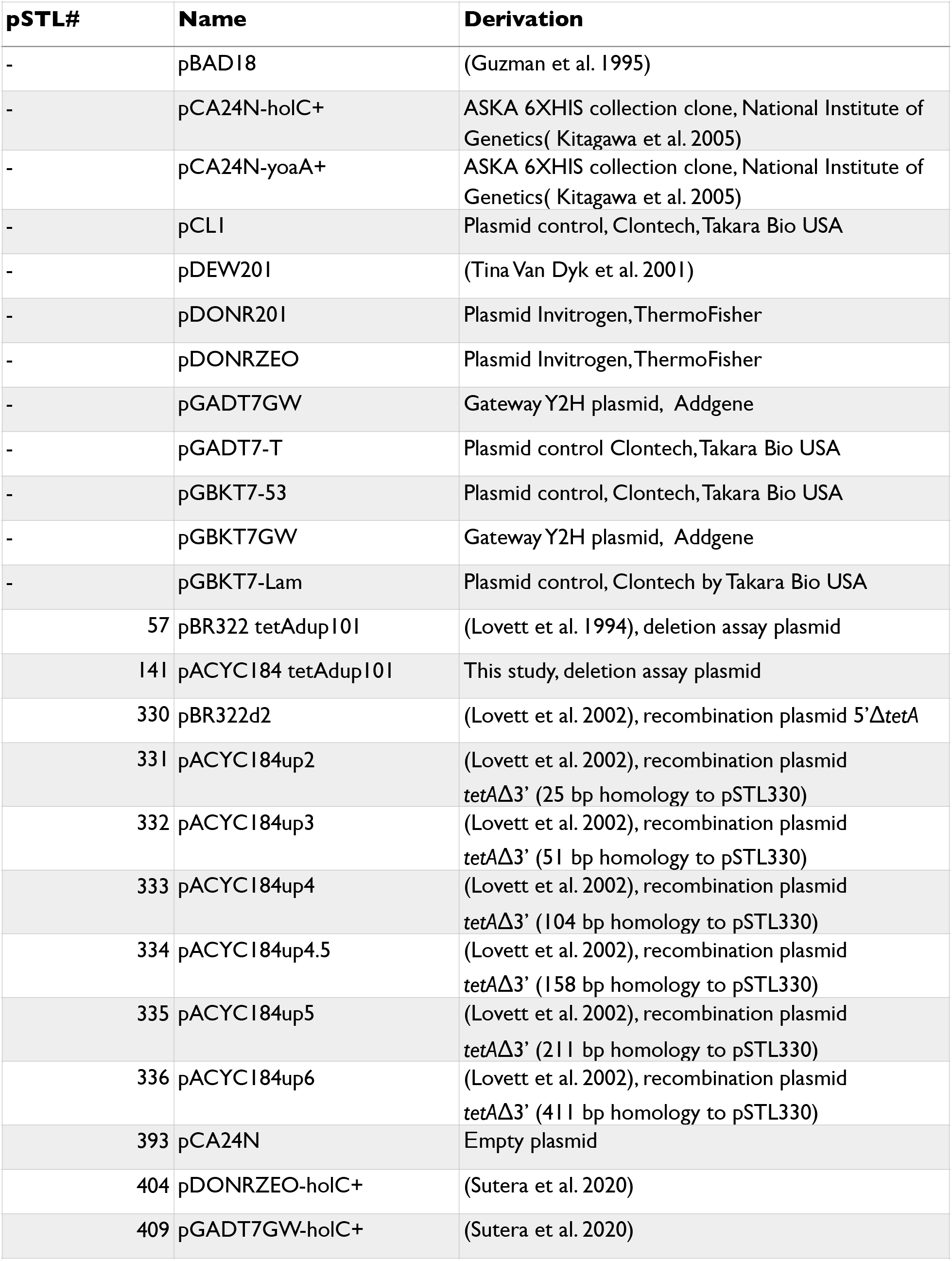
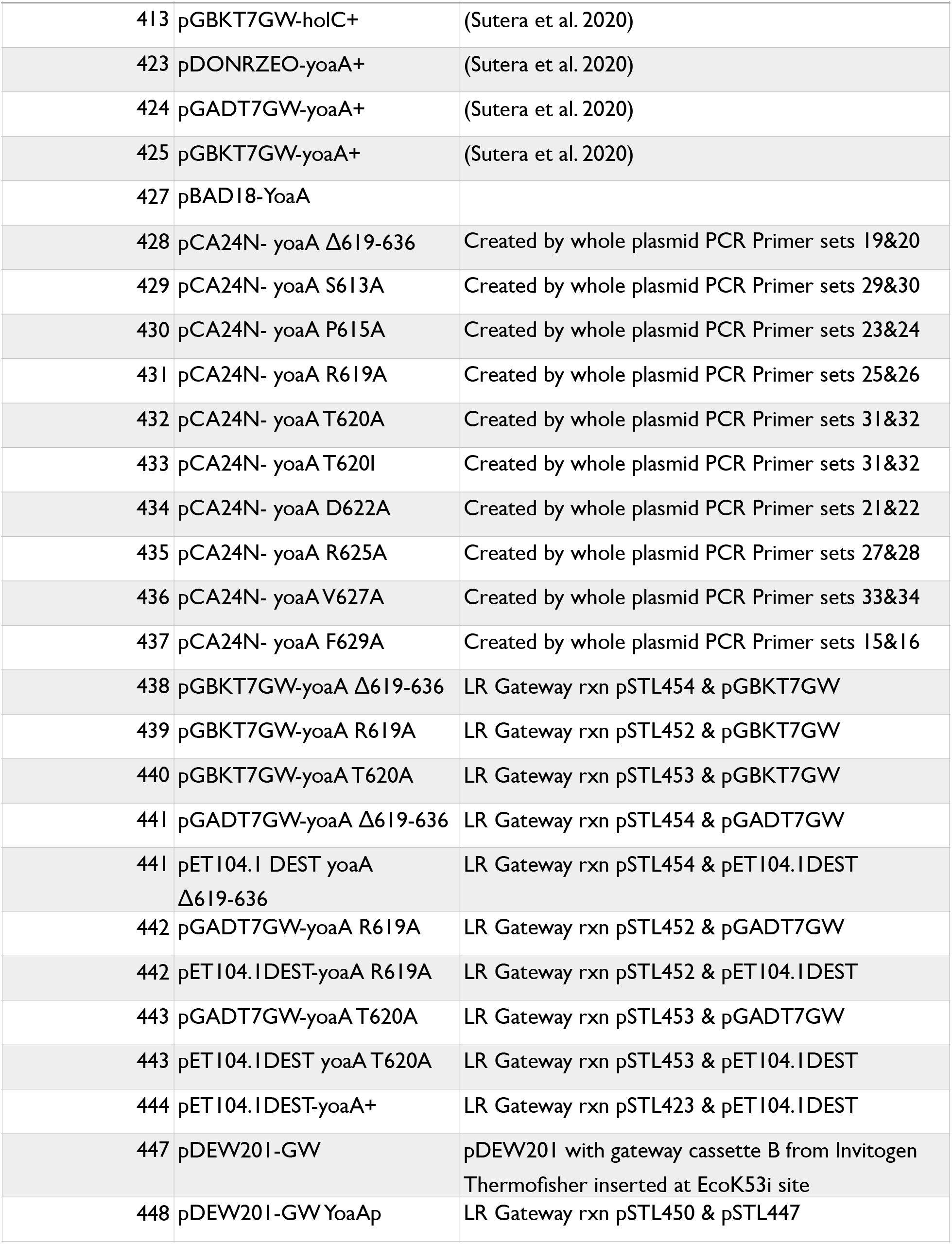
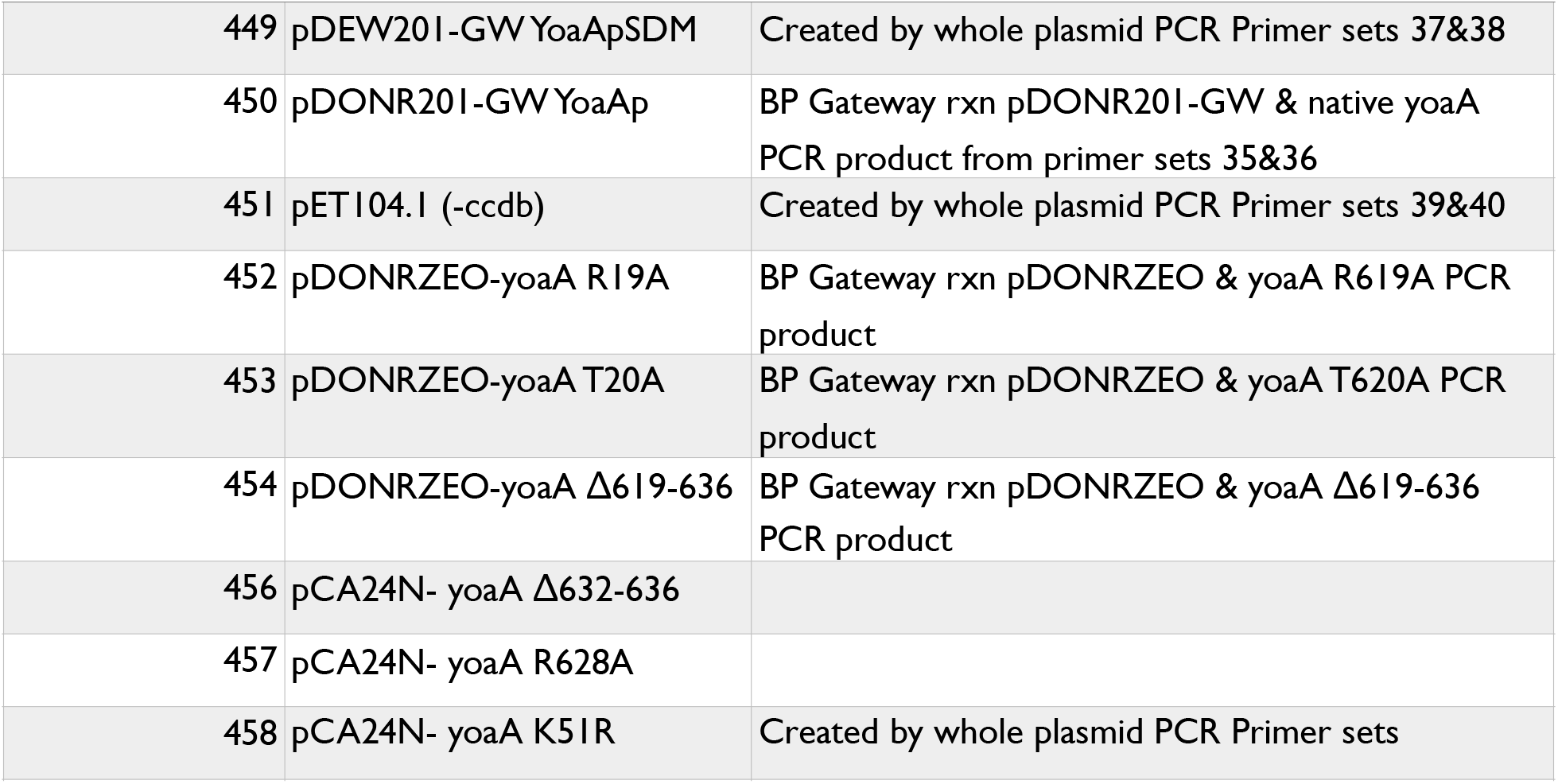
Plasmids.

### Complementation and toxicity assays

Plasmids including high copy number pCA2N vector, pCA2N-YoaA+ (41) and the indicated YoaA site-directed mutants were introduced strain STL9813 (*yoaA*Δ::FRT). Cultures were grown in LB +Cm to log phase, OD600 =0.4 upon which time they were serially diluted into 56/2 buffer and plated on LB medium containing 37.5 ng/ml AZT. For the toxicity assays, MG1655 strains carrying the pCA24N-derived plasmids were inoculated into LB-Ap media in Costar 96 well assay plates and shaken at 37° in a BioTek Cytation plate reader, measuring OD600 every 15 minutes. When cultures reached an OD of 0.2-0.4 IPTG was added to 1/2 the cultures to 1 mM and incubated an additional 3 hours.

### Protein expression and Western blots

BBD-YoaA wildtype (pET104.1DEST-yoaA), BBD-YoaA mutants (pET104.1 yoaA T620A, pET104.1 yoaA T3, pET104.1DEST-yoaA R619A) and the empty pET104.1 vector (listed in Table 1) were expressed from the *E. coli* BL21(DE3) strain. The strains were grown in LB media at 37°C with a final concentration of 100 μg/mL ampicillin, and the cells were induced with 1 mM IPTG for 2 hours. The cells of all overexpressing strains were concentrated by centrifugation at 4700 rpm for 30 minutes at room temperature, concentrated 1:100 in Tris-Sucrose (50 mM Tris-HCl, 10% w/v sucrose, pH 7.5), and stored at -80°C.

Crude cell extracts were prepared with lysozyme lysis using 1µM DTT and 0.1 mg/mL lysozyme (United States Biochemical) in Tris-Sucrose. After a 5-minute incubation on ice, NaCl was added to a final concentration of 0.2 M, and the extract was incubated an additional 25-minute incubation, after which the cells were heat shocked at 37°C for 15 seconds, then transferred to on ice for 30 seconds. Following two heat shocks, the lysed cells were centrifuged, and the crude lysate supernatant was collected. BBD-YoaA protein samples, combined with an equal volume of 2X FSB (0.12 M Tris-HCl pH 6.8, 3.8% SDS, 19% glycerol, 1.43 M beta-mercaptoethanol, 1mg/ml Bromophenol Blue). Samples were subject to polyacrylamide gel electrophoresis in 12% polyacrylamide gels and transferred to PVDF membrane utilizing Bio-rad transblot apparatus at 100 volts and 400 milliamps for 75 minutes. The western blot analysis was performed according to the QIAexpress detection kit and the protocol (Qiagen) with the following modifications: the BBD-YoaA was detected with a dilution of 1:1000 Neutratavidin Antibody (ThermoFisher) in 10% nonfat milk in for 1 hour. Gel was then washed for 4x 10 minutes with TBS-T wash buffer (20mM 1M Tris-HCl pH 7.5, 500 mM NaCl, 0.05% Tween-20, 0.2% Triton X-100). Imaging was performed on a Bio-Rad ChemiDoc system.

### Construction of GAL4 Activation Domain and Binding Domain fusions to HolC and YoaA For Yeast Two-Hybrid System

Bacterial colony PCR with high fidelity DNA polymerase Phusion (New England Biolabs) was used to obtain the wild type alleles of *holC* and *yoaA* for construction of GAL4 fusions for the yeast two hybrid analysis as previously described (16). Gateway cloning technology from Invitrogen® was used to transfer the mutations created in pCA24N-YoaA to the GAL4 activation and binding domain plasmids, pGADT7GW and pGBKT7GW, respectively. Primers were used in a high fidelity PCR reaction using Pfu DNA polymerase obtained from Agilent®. Subsequent PCR reactions were subjected to DpnI to destroy the template plasmid. Following purification of the PCR products (BioBasic kits), *yoaA* mutant fragments (*attB1-yoaA* mutant*– attB2*) were cloned into pDONRZEO using the enzyme BP Clonase II and subsequently into either the pGADT7GW or pGBKT7GW vectors, using LR Clonase II.

### Yeast Two-Hybrid Analysis

We followed procedures and used controls from the Matchmaker Gold Yeast Two-Hybrid System from Takara Bio, with the following modifications. Negative controls consisted of the ones suggested in the Matchmaker Gold Yeast Two-Hybrid System with the addition of *holC* and *yoaA* activation domain hybrid plasmids paired with the binding domain empty plasmid. Similarly *holC* and *yoaA* binding domain plasmids were paired with the activation domain empty plasmid vector. A single colony of all controls and combinations of the activation and binding domain plasmids were grown in 5 mls either leucine or leucine and tryptophan dropout media for 20 hours. Following such incubation cultures were diluted 1:5 and 1:50 in sterile water. 1/20 of this dilution was plated on the following media: YEPD, -leucine, -tryptophan, -leucine and tryptophan, -histidine and -adenine. YEPD is the universal medium for which all cultures will grown. The plates that lack either leucine, tryptophan or both leucine and tryptophan are controls to test plasmid retention. The histidine deficient plates test for lower stringency protein-protein interactions while adenine deficient plates test for a higher stringency. Plates were incubated for two or three days at 30°C.

### Luciferase gene expression assays

Luciferase fusion construct plasmids were based on plasmid pDEW201 (42) with an inserted GATEWAY-cloning *attR* site-specific recombination cassette. Plasmid construction was completed using GATEWAY cloning (Life Technologies) from PCR amplified products with the primer pair yoaApromoterGWF 5’-ggggacaagtttgt acaaaaaagc aggcttcCATTTTGTCCTCA TTATACTTCC AT-3’ and yoaApromoterGWR 5’-ggggaccact ttgtacaaga aagctgggtc ACTACCCCCT GTTGATTTGA ACAGG-3’. Products were recombined with the BP reaction into the pDONR201 GATEWAY plasmid vector. Verified pDONR201GW-yoaAp were recombined using the LR reaction into the GATEWAY pDEW201 Ap LuxCDABE vector. To create the yoaApSDM with the mutated LexA box, site directed mutagenesis (Quikchange Agilent Technologies) on the pDEW201GW-yoaAP was conducted according to the manufacturer’s instructions, using overlapping PCR primers with the target mutation, yoaALexAp1 5’-GCGCCCTCAT CCTGACATAA TGTCCCTTCA AATCAAGGGA CGGTAGTGTG ACGGAC-3’ and yoaALexAp2 5’-GTCCGTCACA CTACCGTCCC TTGATTTGAA GGGACATTAT GTCAGGATGA GGGCGC-3’ and amplification with the Phusion High-Fidelity DNA Polymerase PCR kit (New England BioLabs). Constructs were sequence verified.

Luminescence and OD600 were measured using BioTek Cytation 1 Plate Reader and Costar 96 Well Assay Plate (treated polystyrene, black plate, clear bottom). Colonies were inoculated in LB media in tubes, shaking until OD_600_ = 0.5 after which they were diluted 1:100 in LB and grown again to ensure log phase growth. In the 96 well plates, cells were diluted 1:100 and grown for 2 hours, before being treated with 1.25 ng/mL AZT. Bioluminescence was measured and normalized to the OD_600_ yielding relative luminescence units (RLU) every 15 minutes and data are averages of 4 independent replicate cultures.

### Recombination, mutagenesis and deletion assays

Recombination assays were performed as previously described (26), with crossovers between plasmids carrying various homologies, selected by Tc-resistance. Wt (MG1655) and *yoaA* (STL9813) mutant strains were assayed in parallel on multiple days. Transversion mutagenesis assays used previously described alleles of chromosomal *lacZ* that revert to Lac+ only by a A- to T-transversion (33). Cultures grown in LB, then were split with expression induced in one by addition of 0.2% arabinose. Cultures were grown for an additional 2 h and then diluted and plated on minimal lactose media to count Lac+ revertants and LB to count the total number of cells in the culture. Plasmids pSTL57 (Ap^r,^ ColE1 replicon (43)) and pSTL141 (Cm^r^, p15a replicon) were used to measure deletion formation between 101 bp repeats in *tetA* as previously described. (43) For experiments with induced expression of YoaA, pBAD18-YoaA or pBAD18 (Ap^r^, ColE1 (44)) were introduced in strains carrying the pSTL141 deletion reporter. Cultures grown in LB Ap Cm were split with expression induced in one by addition of 0.2% arabinose. Cultures were grown for an additional 2 h and then diluted and plated on LB Ap Cm Tc.

---

**Table 2.**
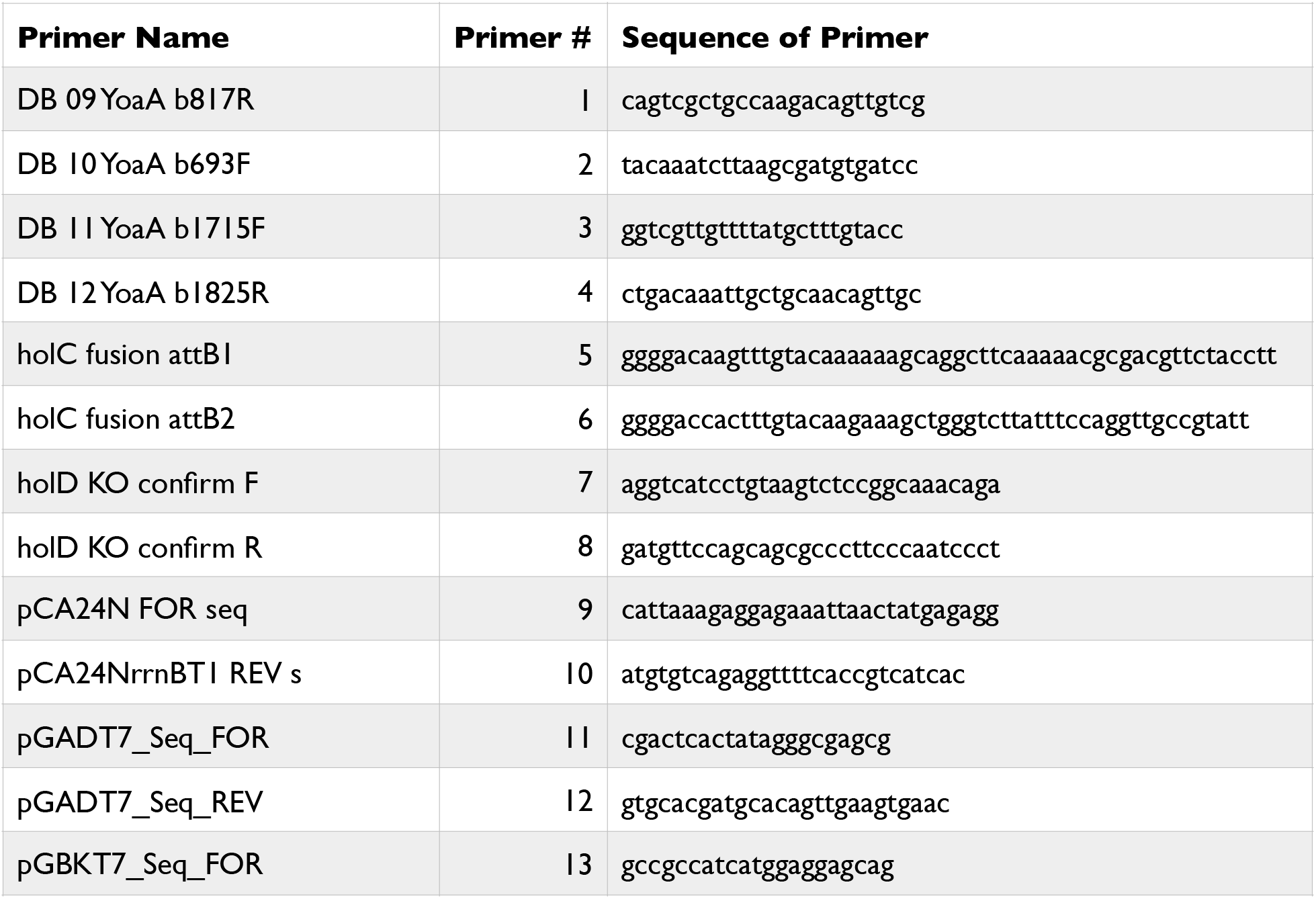
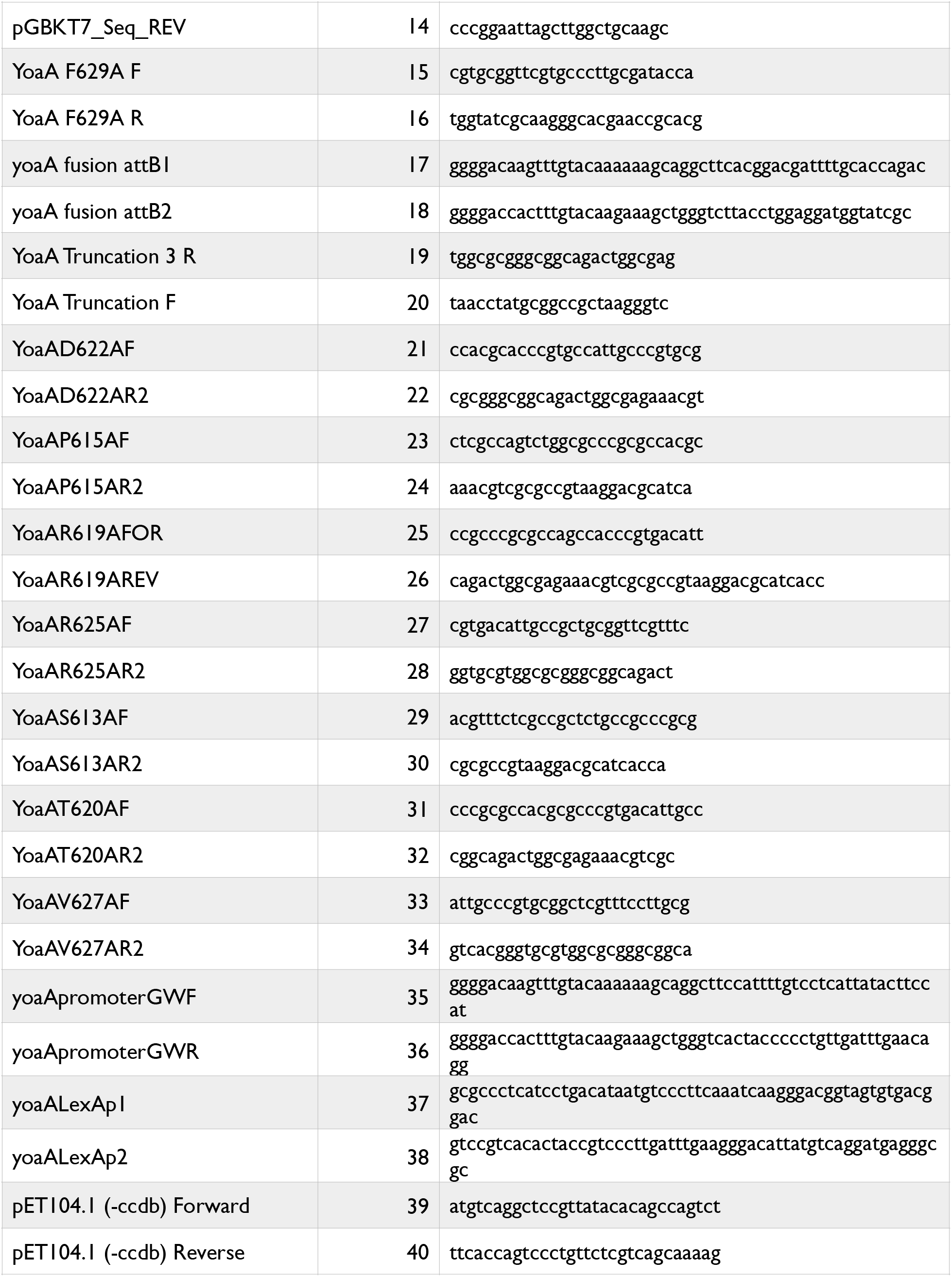
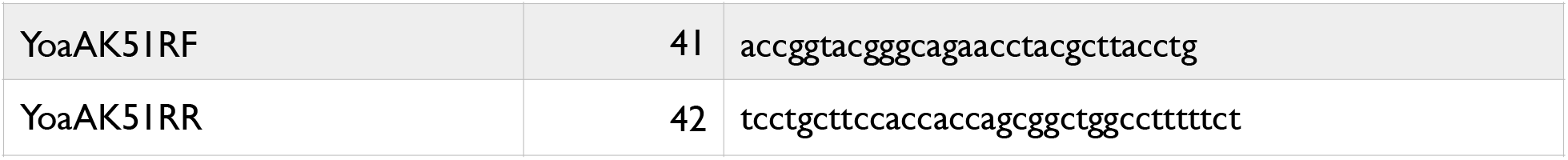
Primers.

## Acknowledgments

This work was support by NIGMS R01 grant GM51753 to STL and T32 GM007122 to THS. We thank Tracey Seier for initial deletions assays of *yoaA* and *dinG* strains, Deani Cooper and Laura Brown for initial work on the expression of YoaA and Kyle McSweeney and McKay Shaw for construction of some of the site-directed mutants.

---

**Supplemental Figure 1.**
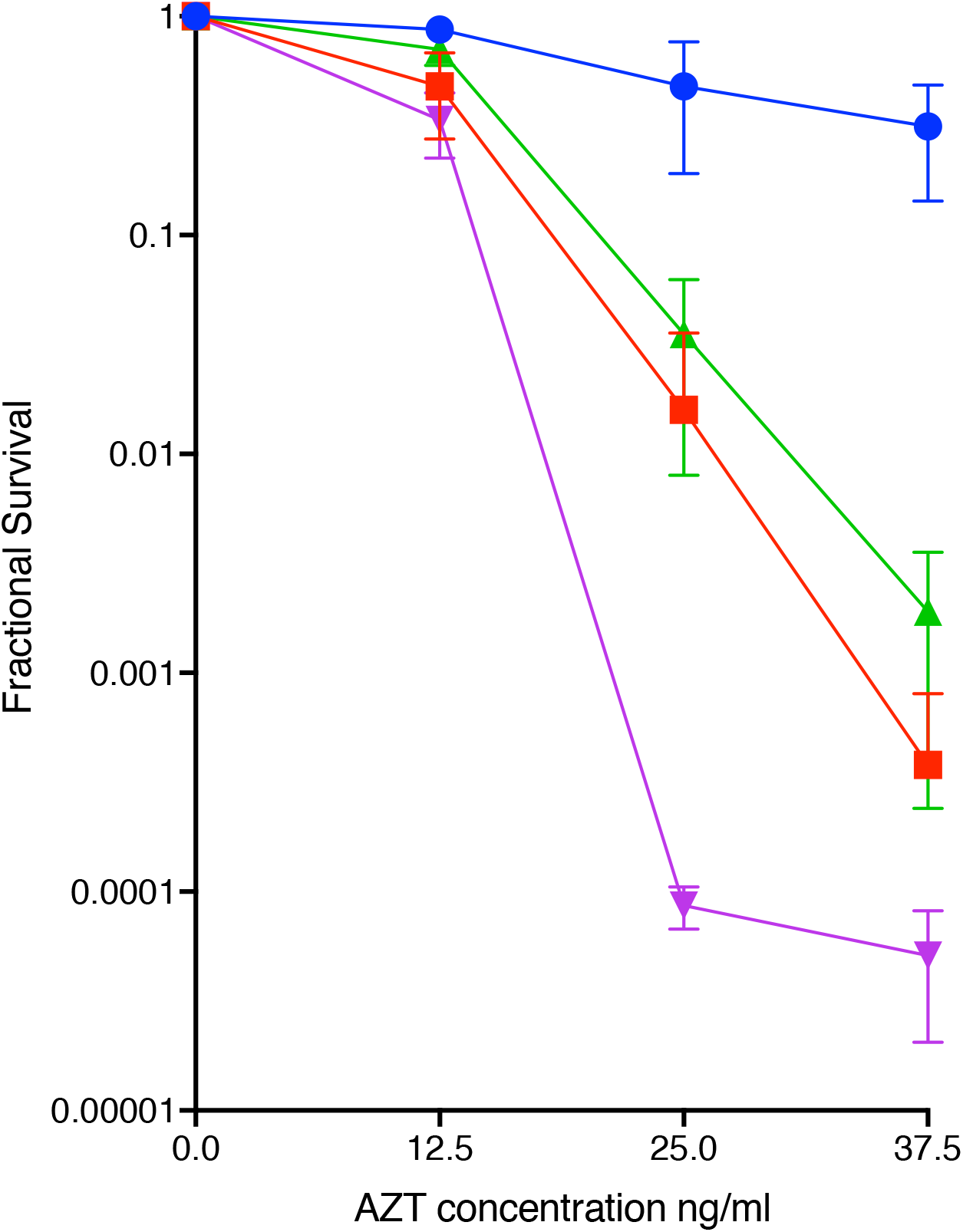
AZT survival of wt (blue circles), or *yoaA*Δ619-636 (purple triangles), *yoaA* R619A (red squares) *yoaA* T720A (green triangles) mutants at the natural *yoaA* chromosomal locus.

## Notes

### Competing Interest Statement

The authors have declared no competing interest.

### Summary of Updates

New data added to the revision: Fig. 5 toxicity of YoaA is dependent on the C-terminal domain; Fig 7 YoaA effects on genome stability now includes data on dinG and yoaA effects on tandem repeat deletion and yoaA effects on A to T transversion mutations in wt and ndk mutants, Supplemental Figure 1 AZT sensitivity of yoaA interaction mutants transferred to the chromosome New model figure, Fig. 8 The text has been extensively revised to improve the narrative and comment on the new data.

## Bibliography

1. Lehmann AR. 2003. DNA repair-deficient diseases, xeroderma pigmentosum, Cockayne syndrome and trichothiodystrophy. Biochimie 85:1101–11.

2. Wu Y, Suhasini AN, Brosh RM. 2009. Welcome the family of FANCJ-like helicases to the block of genome stability maintenance proteins. Cellular and molecular life sciences : CMLS 66:1209–1222.

3. Bharti SK, Awate S, Banerjee T, Brosh RM. 2016. Getting Ready for the Dance: FANCJ Irons Out DNA Wrinkles. Genes (Basel) 7.

4. Abe T, Ooka M, Kawasumi R, Miyata K, Takata M, Hirota K, Branzei D. 2018. Warsaw breakage syndrome DDX11 helicase acts jointly with RAD17 in the repair of bulky lesions and replication through abasic sites. Proc Natl Acad Sci U S A 115:8412–8417.

5. Lewis LK, Jenkins ME, Mount DW. 1992. Isolation of DNA damage-inducible promoters in Escherichia coli: regulation of polB (dinA), dinG, and dinH by LexA repressor. Journal of Bacteriology 174:3377–3385.

6. Lewis LK, Mount DW. 1992. Interaction of LexA repressor with the asymmetric dinG operator and complete nucleotide sequence of the gene. Journal of Bacteriology 174:5110–5116.

7. Courcelle J, Khodursky A, Peter B, Brown PO, Hanawalt PC. 2001. Comparative gene expression profiles following UV exposure in wild-type and SOS-deficientEscherichia coli. Genetics 158:41–64.

8. Voloshin ON, Vanevski F, Khil PP, Camerini-Otero RD. 2003. Characterization of the DNA damage-inducible helicase DinG from Escherichia coli. The Journal of biological chemistry 278:28284–28293.

9. Voloshin ON, Camerini-Otero RD. 2007. The DinG protein from Escherichia coli is a structure-specific helicase. The Journal of biological chemistry 282:18437–18447.

10. Thakur RS, Desingu A, Basavaraju S, Subramanya S, Rao DN, Nagaraju G. 2014. Mycobacterium tuberculosis DinG is a structure-specific helicase that unwinds G4 DNA: implications for targeting G4 DNA as a novel therapeutic approach. J Biol Chem 289:25112–36.

11. Boubakri H, de Septenville AL, Viguera E, Michel B. 2010. The helicases DinG, Rep and UvrD cooperate to promote replication across transcription units in vivo. EMBO J 29:145–157.

12. Brown LT, Sutera VA, Jr., Zhou S, Weitzel CS, Cheng Y, Lovett ST. 2015. Connecting Replication and Repair: YoaA, a Helicase-Related Protein, Promotes Azidothymidine Tolerance through Association with Chi, an Accessory Clamp Loader Protein. PLoS Genet 11:e1005651.

13. Cooper DL, Lovett ST. 2011. Toxicity and tolerance mechanisms for azidothymidine, a replication gap-promoting agent, in Escherichia coli. DNA Repair 10:260–270.

14. Watanabe K, Tominaga K, Kitamura M, Kato JI. 2016. Systematic identification of synthetic lethal mutations with reduced-genome Escherichia coli: synthetic genetic interactions among yoaA, xthA and holC related to survival from MMS exposure. Genes Genet Syst 91:183–188.

15. Butland G, Peregrin-Alvarez J, Li J, Yang W, Yang X, Canadien V, Starostine A, Richards D, Beattie B, Krogan N, Davey M, Parkinson J, Greenblatt J, Emili A. 2005. Interaction network containing conserved and essential protein complexes in Escherichia coli. Nature 433:531–537.

16. Sutera VA, Weeks SJ, Dudenhausen EE, Rappe Baggett HB, Shaw MC, Brand KA, Glass DJ, Bloom LB, Lovett ST. 2020. Alternative complexes formed by the Escherichia coli clamp loader accessory protein HolC (x) with replication protein HolD (ψ) and repair protein YoaA. DNA Repair in press.

17. Reyes-Lamothe R, Sherratt DJ, Leake MC. 2010. Stoichiometry and architecture of active DNA replication machinery inEscherichia coli. Science 328:498–501.

18. McHenry CS. 2011. DNA replicases from a bacterial perspective. Annu Rev Biochem 80:403–36.

19. Kelman Z, Yuzhakov A, Andjelkovic J, O’Donnell M. 1998. Devoted to the lagging strand-the subunit of DNA polymerase III holoenzyme contacts SSB to promote processive elongation and sliding clamp assembly. EMBO J 17:2436–49.

20. Xiao H, Dong Z, O’Donnell M. 1993. DNA polymerase III accessory proteins. IV. Characterization of chi and psi. J Biol Chem 268:11779–84.

21. Gao D, McHenry CS. 2001. Tau binds and organizes Escherichia coli replication proteins through distinct domains. Domain III, shared by gamma and tau, binds delta delta ‘ and chi psi. J Biol Chem 276:4447–53.

22. Sutera VA, Weeks SJ, Dudenhausen EE, Baggett HBR, Shaw MC, Brand KA, Glass DJ, Bloom LB, Lovett ST. 2021. Alternative complexes formed by the Escherichia coli clamp loader accessory protein HolC (x) with replication protein HolD (psi) and repair protein YoaA. DNA Repair (Amst) 100:103006.

23. Gulbis JM, Kazmirski SL, Finkelstein J, Kelman Z, O’Donnell M, Kuriyan J. 2004. Crystal structure of the chi:psi sub-assembly of the Escherichia coli DNA polymerase clamp-loader complex. Euro J Biochem 271:439–449.

24. Cheng K, Wigley DB. 2018. DNA translocation mechanism of an XPD family helicase. Elife 7.

25. Duigou S, Silvain M, Viguera E, Michel B. 2014. ssbgene duplication restores the viability of ΔholCand ΔholDEscherichia colimutants. PLoS Genet 10:e1004719.

26. Lovett ST, Hurley RL, Sutera VA, Jr., Aubuchon RH, Lebedeva MA. 2002. Crossing over between regions of limited homology in Escherichia coli. RecA-dependent and RecA-independent pathways. Genetics 160:851–9.

27. Dutra BE, Sutera VA, Jr., Lovett ST. 2007. RecA-independent recombination is efficient but limited by exonucleases. Proc Natl Acad Sci U S A 104:216–21.

28. Persky NS, Lovett ST. 2008. Mechanisms of recombination: lessons from E. coli. Crit Rev Biochem Mol Biol 43:347–70.

29. Lovett ST. 2017. Template-switching during replication fork repair in bacteria. DNA Repair (Amst) 56:118–128.

30. Lu Q, Zhang X, Almaula N, Mathews CK, Inouye M. 1995. The gene for nucleoside diphosphate kinase functions as a mutator gene in Escherichia coli. J Mol Biol 254:337–41.

31. Miller JH, Funchain P, Clendenin W, Huang T, Nguyen A, Wolff E, Yeung A, Chiang JH, Garibyan L, Slupska MM, Yang H. 2002. Escherichia coli strains (ndk) lacking nucleoside diphosphate kinase are powerful mutators for base substitutions and frameshifts in mismatch-repair-deficient strains. Genetics 162:5–13.

32. Schaaper RM, Mathews CK. 2013. Mutational consequences of dNTP pool imbalances in E. coli. DNA Repair (Amst) 12:73–9.

33. Seier T, Padgett DR, Zilberberg G, Sutera VA, Jr., Toha N, Lovett ST. 2011. Insights into mutagenesis using Escherichia coli chromosomal lacZ strains that enable detection of a wide spectrum of mutational events. Genetics 188:247–62.

34. Cooper DL, Harada T, Tamazi S, Ferrazzoli AE, Lovett ST. 2021. The role of replication clamp-loader protein HolC of Escherichia coli in overcoming replication /transcription conflicts. mBio in press.

35. Little JW, Edmiston SH, Pacelli LZ, Mount DW. 1980. Cleavage of the Escherichia coli lexA protein by the recA protease. Proc Natl Acad Sci U S A 77:3225–9.

36. Lovett ST, Drapkin PT, Sutera VA, Jr., Gluckman-Peskind TJ. 1993. A sister-strand exchange mechanism for recA-independent deletion of repeated DNA sequences in Escherichia coli. Genetics 135:631–42.

37. Miller JH. 1992. A Short Course in Bacterial Genetics. Cold Spring Harbor Press,New York.

38. Bernard MA, Ray NB, Olcott MC, Hendricks SP, Mathews CK. 2000. Metabolic functions of microbial nucleoside diphosphate kinases. J Bioenerg Biomembr 32:259–67.

39. Sherman F, Fink G, Hicks J. 1987. Methods in Yeast Genetics: a Laboratory Course Manual. Cold Spring Harbor Press, Cold Spring Harbor, NY.

40. Dower W, Miller J, Ragsdale C. 1988. High efficiency transformation ofE. Coliby high voltage electroporation. Nucleic Acids Research 16:6127–6145.

41. Kitagawa M, Ara T, Arifuzzaman M, Ioka-Nakamichi T, Inamoto E, Toyonaga H, Mori H. 2005. Complete set of ORF clones of Escherichia coli ASKA library (a complete set of E. coli K-12 ORF archive): unique resources for biological research. DNA research : an international journal for rapid publication of reports on genes and genomes 12:291–299.

42. Van Dyk TK, DeRose EJ, Gonye GE. 2001. LuxArray, a high-density, genomewide transcription analysis ofEscherichia coliusing bioluminescent reporter strains. J Bacteriol 183:5496–505.

43. Lovett ST, Gluckman TJ, Simon PJ, Sutera VA, Jr., Drapkin PT. 1994. Recombination between repeats inEscherichia coliby arecA-independent, proximity-sensitive mechanism. Mol Gen Genet 245:294–300.

44. Guzman L, Belin D, Carson M, Beckwith J. 1995. Tight regulation, modulation, and high-level expression by vectors containing the arabinose PBAD promoter. J Bacteriol 177:4121–30.

